# Effects of Turbidity and Habitat Complexity on the Foraging Behavior of the Black Bullhead (*Ameiurus melas*)

**DOI:** 10.1101/2024.02.12.579876

**Authors:** Bálint Preiszner, Anna Szolnoki, István Czeglédi, Tibor Erős

## Abstract

The role of vertebrate scavengers in nutrient flow of aquatic ecosystems has recently been emphasized. Evidence is accumulating that scavenging is an alternative tactic for certain predatory fish, but their foraging choices between scavenging and predation under varying environmental conditions have not been studied. In this experiment, we presented black bullheads, *Ameiurus melas* (Rafinesque, 1820), with both live and carcass prey of tubenose gobies, *Proterorhinus semilunaris* (Heckel, 1837), in aquaria with varying levels of turbidity and habitat complexity. Our findings revealed that black bullheads exhibit higher consumption of carcass prey in complex habitat and clear water condition. We also provide evidence of the active predatory behavior displayed by black bullheads, as a greater consumption of live prey was observed in simple habitat and turbid water. We did not find any relationship between individual exploratory behavior and the choice between carcass and live prey in black bullheads. Our results suggest that the choice between scavenging versus predatory foraging strategy under varying environmental conditions is influenced by the behavior of both consumer and prey. The ability of black bullheads to adapt their foraging strategy in different environments may contribute to their widespread success as an invasive species.

## Introduction

Trophic interactions are one of the key organizing principles within ecosystems, which define the ecological role of species within communities (Bump, Peterson, & Vucetich, 2009; Hooper et al., 2005). Trophic relationships were traditionally viewed as relatively constant, at least in most food web models (Allesina, Alonso, & Pascual, 2008). However, due to a series of recent research on individual and species-level flexibility of foraging behavior, these static models should be reconsidered. For example, it has been shown that sometimes vertebrate species that used to be regarded as predatory tend to consume carcasses more often than previously thought (DeVault, Rhodes, & Shivik, 2003), rendering their role more complex in trophic relationships and nutrient recycling (Boros, Czeglédi, Erős, & Preiszner, 2020). This increased awareness of vertebrate scavenging behavior sparked a recrudescence in scavenging research (Amorós, Gil-Sánchez, López-Pastor, & Moleón, 2020; Benbow, Tomberlin, & Tarone, 2015; Kane, Healy, Guillerme, Ruxton, & Jackson, 2017; Ogada, Torchin, Kinnaird, & Ezenwa, 2012). However, most studies focused on terrestrial species (Beasley, Olson, & Devault, 2012; Olea, Mateo-Tomás, & Sánchez-Zapata, 2019) and less is known about aquatic vertebrates in this regard, perhaps due to the technological difficulties of underwater studies (Boros et al., 2020; but see Polačik, Jurajda, Blažek, & Janáč, 2015; Preiszner et al., 2020).

One of the key concepts of foraging theory is that behavioral flexibility, i.e. the ability of an individual to change its existing behavioral patterns (West-Eberhard, 1989), is a fundamental tool for every forager that allows to adapt its tactics and diet choice according to altering conditions (Dill, 1983; Sih & Christensen, 2001). Environmental variability may induce alteration of activity budgets or foraging habitat choice as it was shown for a variety of taxa (Aublet, Festa-Bianchet, Bergero, & Bassano, 2009; see e.g. Litzow & Piatt, 2003; Noreika, Bartomeus, Winsa, Bommarco, & Öckinger, 2019). However, in many cases foraging flexibility assumes a plasticity in food types consumed. For example, fish are known to shift between zooplankton and benthic macroinvertebrate food items, contributing to their invasion success (Hayden et al., 2014). Therefore, such flexibilities in foraging behavior may alter fundamental ecological processes, underlining their importance (Abrams, 2010).

Foraging interactions between consumers and their food items correlate with the parties’ behavior. It has been proposed that some proximate mechanisms underlying this relationship are mediated directly or indirectly by environmental conditions. In aquatic environment, behavior related to trophic interactions is known to vary along gradients of environmental factors such as temperature, turbidity, and habitat complexity. Suspended material in the water for example, may have several physiological effects on aquatic organisms, which in turn may influence their behavior, and thus their trophic relationships (Henley, Patterson, Neves, & Dennis Lemly, 2000). A change in turbidity may trigger a shift in habitat use; for example, juvenile bluegills, *Lepomis macrochirus* (Rafinesque, 1819), move offshore from nearshore habitats when met turbid conditions, and at the same time, their reaction distance is inversely related to turbidity (Miner & Stein, 1996). Multipredator interactions are also known to vary along the turbidity gradient, although with contrasting outcomes, i.e. turbidity may increase, decrease or may not affect predator efficiency (VanLandeghem, Carey, & Wahl, 2011). Altered predation rate may in turn facilitate adjustment of anti-predator behavior of prey, as shown by Lehtiniemi *et al*. (2005) in pike, *Esox lucius* (Linnaeus, 1758). Another environmental factor, habitat complexity affects foraging efficiency for example by altering activity, searching and handling time (Anderson, 1984; Murray, Stillman, & Britton, 2016; Priyadarshana, Asaeda, & Manatunge, 2001). Spatial complexity may influence feeding behavior differently in accordance with foraging strategy; for example it may influence feeding success through shaping prey behavior, but it may also regulate the occurrence of ambush versus pursuit predator strategy (Flynn & Ritz, 1999; Říha et al., 2021). Habitat heterogeneity is also known to mediate trophic interactions by influencing intraspecific variation of trophic niche breadths and interspecific niche overlaps (Dias, Tófoli, DaSilva, Gomes, & Agostinho, 2022), promoting coexistence in high-complexity habitats. Such environmental factors may also have an impact on proportion of scavenging within carnivores’ diet, the mechanisms of which yet little is known. Understanding the effects of environmental conditions on foraging decisions is especially relevant in ecosystems where spatial and temporal alteration of such conditions are recurring.

Personality plays an important role in shaping the behavior of individuals, which in turn affects many aspects of their lives. For example, bold individuals may be more successful competitors in some context (Ericsson et al., 2021), but shy individuals can have decreased predation risk (Miyamoto & Araki, 2020). A relatively easily quantifiable personality trait, the explorative behavior has been studied across a variety of taxa in various contexts (Reader, 2015). It may have positive and negative effects on foraging behavior, and the outcome can be context dependent. Therefore, the trade-off between costs and benefits of explorative behavior may impact the foraging tactic and thus ultimately the overall foraging success of an individual in different ecological contexts.

Here we studied the effects of environmental conditions on the food choice of black bullheads, *Ameiurus melas* (Rafinesque, 1820), in aquaria. This North American species had been introduced to Europe in the nineteenth and twentieth centuries and became one of the most invasive species in many countries (Copp et al., 2016), including Hungary (Ferincz et al., 2016; Takács, Abonyi, Bánó, & Erős, 2021; Takács et al., 2017). Although several studies report on its diet (Leunda et al., 2008; Ruiz-Navarro, Britton, Jackson, Davies, & Sheath, 2015), several aspects of its feeding ecology remains unknown, for example how environmental conditions influence the role of scavenging vs predation in its trophic relationships with native and non-native fishes.

Consequently, we offered them tubenose gobies, *Proterorhinus semilunaris* (Heckel, 1837), alive and as fresh carcass simultaneously under different levels of turbidity and habitat complexity. Both species are bottom dwellers, and can be found in the same freshwater ecosystems across Central Europe (Kováč, 2015; Musil, Jurajda, Adámek, Horký, & Slavík, 2010) including Lake Balaton, the largest shallow lake in Central-Europe, where they co-occur in the same habitat (Czeglédi et al., 2019), indicating a high likelihood of trophic interactions between them. Scavenging along with predatory behavior of black bullheads has been suspected in natural environments (see e.g. Leunda et al., 2008; and Snow, Porta, & Robison, 2017), and both has been proved experimentally (scavenging by Preiszner et al., 2020; preying on larval fish by Silbernagel & Sorensen, 2013). Consequently, we hypothesized that live prey and carcass have different functional value for this omnivore fish that varies according to the characteristics of the environment, and which in turn is mirrored by the consumer’s food choice. Specifically, we addressed the following questions: (1) do black bullheads show food choice preference for carcass or live prey? (2) Does food choice vary depending on complexity of the habitat and turbidity of the water? (3) Is explorative behavior related to food choice preference?

## Methods

All animals used in the experiments were captured in the littoral zone of Lake Balaton, Hungary. Black bullheads were collected with fyke nets (net length: 80 cm; moderately expandable throat size: 15 cm; mesh size: 0.8 cm) and were placed in indoor aquaria as soon as possible (maximum three hours after capture). Size range of the selected black bullheads was narrow and varied between 146- and 156-mm standard length (SL – Table 1.). Tubenose gobies were captured with an electrofishing device (IG200/2B, PDC, 50–100 Hz, 350–650 V, max. 10 kW; Hans Grassl GmbH, Germany). Size of the selected tubenose gobies varied between 25- and 38-mm SL (Table 1.), and were small enough to exclude gape size limitation of the consumer. Tubenose gobies chosen as designated live preys were stocked in indoor aquaria as soon as possible (maximum two hours after capture), whereas fresh carcasses were stored in a freezer for a maximum of one month, from where they were removed one hour prior placing them in the experimental tanks.

**Table 1.** Body measurements of the animals immediately before the binary food choice test (mean ± standard deviation): initial body weight (g), SL (standard length - distance between the tip of the head and the base of the caudal fin, mm), maximum height and width of the mouth gape (mm)

|  | **Initial body**<br>**weight (g)** | **SL (mm)** | **Mouth gape**<br>**height (mm)** | **Mouth gape**<br>**width (mm)** |
| --- | --- | --- | --- | --- |
| **Black bullhead**<br>**(N=24)** | 71.63 ± 7.57 | 150.17 ± 2.57 | 22.50 ± 0.93 | 31.17 ± 1.24 |
| **Tubenose goby**<br>**(N=144)** | 0.68 ± 0.18 | 31.71 ± 2.63 | — | — |

During the acclimatization period individuals were placed in holding tanks (length × width × height: 100 × 50 × 50 cm, filled with 200 liters of water) in single species groups. Black bullheads were fed with sinking pellets developed for catfish, while gobies with frozen mosquito (Chironomidae) larvae every other day. Feeding of black bullheads was ceased 48 hours prior to the beginning of the experiments to ensure complete emptying of the intestinal tract.

### Experimental Design

To measure individual exploratory behavior of black bullheads open field tests (Gould, Dao, & Kovacsics, 2009) were carried out. To that end, fish were individually placed in visually separated aquaria (50 × 50 × 40 cm, filled with 60 liters of water), without shelter and aerator. The movement of each individual was recorded from above with Techson TCI MA0 C602 IR A VF IP cameras for 60 minutes (for a sample footage see Online Resource Video S1.). Measurement of body weight, body length and mouth size were taken after the open field tests to minimize stress. Subsequently, black bullheads were individually transferred to randomly chosen experimental tanks (100 × 50 × 50 cm, filled with 120 liters of water). Visual separation between the tanks was applied. Binary food choice tests were started after a 24 h acclimatization period, by introducing three living individuals and three carcasses of tubenose gobies, after measuring their body weight, and body length. In this test, environmental variability was introduced in a 2×2 factorial design by setting up the water turbidity and habitat complexity (Fig. 1.). The water was either clear or turbid (mean ± SD: 0.65 ± 0.24, and 71.70 ± 9.51 Formazin Nephelometric Unit value, respectively), whereas the habitat was either simple (bare bottom) or complex (three equally spaced stones as shelters, each approximately 15×15 cm in area, and similar in volume). Each treatment combination was repeated 6 times, thus a total of 24 repetitions were performed. Black bullheads were allowed to feed for 24 hours, after which the test was terminated. The remaining gobies were counted, and additionally, any sign of partial consumption was also recorded.

**Fig. 1.**
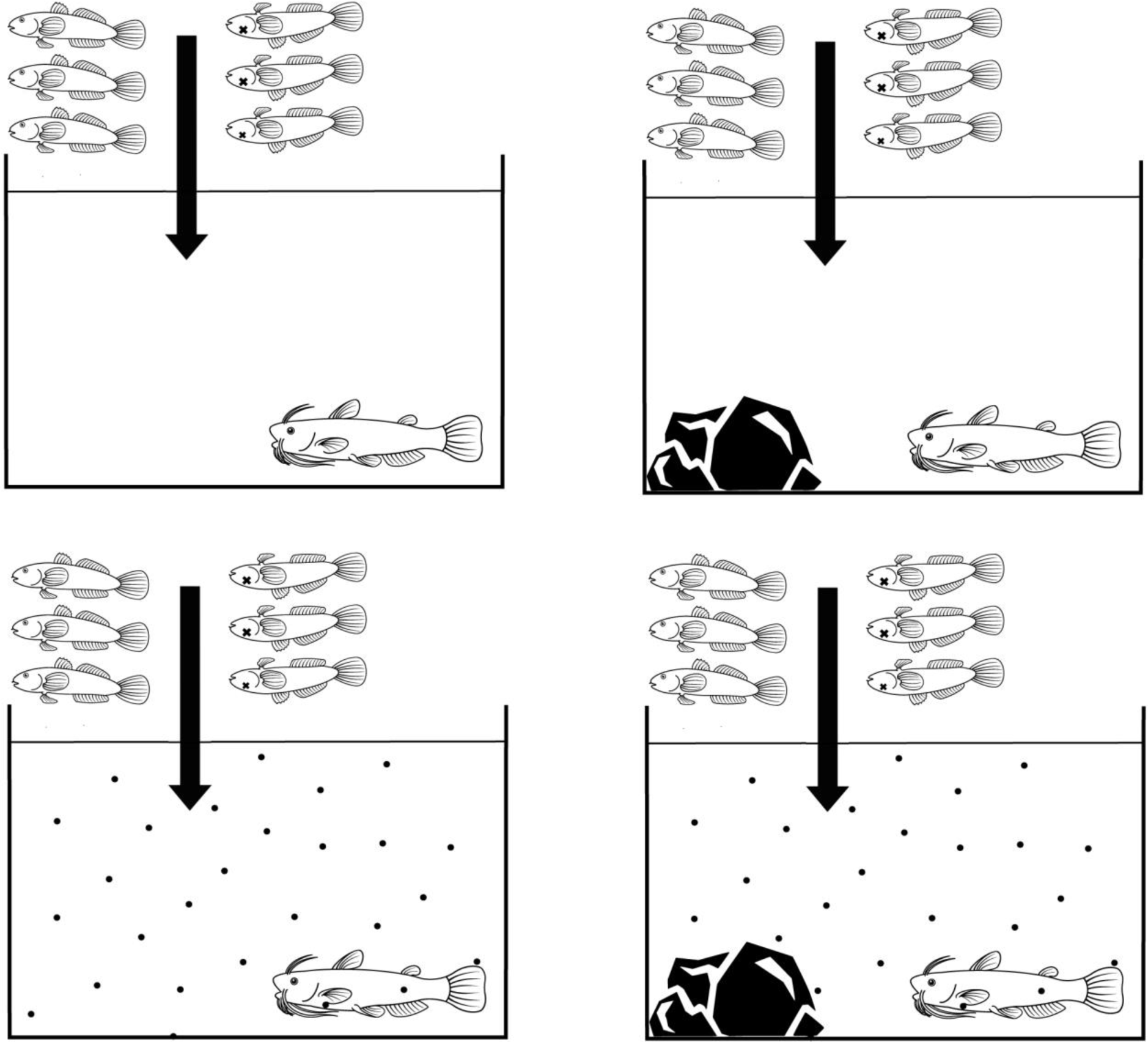
Experimental design for evaluating prey choice of black bullheads (for details, see the Methods section)

A constant temperature of 19 °C and a 12:12 h light:dark period were maintained during the captivity and experiments. The aquarium water consisted of a 50-50% mixture of aged tap water and filtered, sedimented water from Lake Balaton. Aeration was provided in the holding and experimental tanks continuously, except for the duration of exploration tests.

### Statistical Analysis

Movement of black bullhead individuals was analyzed by the ToxTrack software (Rodriguez et al., 2018) from the video recordings. To assess exploration of the arena two arena sizes were used; the full arena, and the actual open field, i.e. the center of the arena. Centre was defined as the area at least 7 cm from the walls, which corresponds for 52% of the full experimental arena area. For software analysis, the experimental arena was divided into regular square-shaped, non-overlapping units of the same size (Online Resource Fig. S1., panel a.). The exploration rate is equal to the ratio of the number of explored units and the total number of units within the experimental arena. Explored units correspond to those within which the software detected the tested individual during the analysis. Since exploration rate in the full arena and exploration rate in the center strongly correlated (r_s_ = 0.973, p < 0.001), the data for the full arena was used as exploration rate in further analysis. Time spent in the center of the arena (henceforth “center time”), was calculated proportionally to total experiment time. Mobility rate is the time spent moving (above the threshold speed of 1 mm/s) by the individual divided by the length of the experiment.

To assess carcass preference, we calculated Ivlev’s electivity index (*E_i_*) based on Ivlev and Lechowicz (1961; and 1982, respectively) using the following formula:

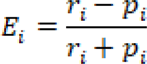

Where *r_i_* is the relative consumption of food type *i* and *p_i_* is the relative availability of that food type in the environment. The index can take a value between plus and minus one. Zero means random choice, where the proportion of consumed food types is equal to their availability in the environment. Positive and negative values represent higher and lower proportion of consumption of the reference food type relative to its proportion in available food items. In our study, we expressed the consumption rate as a preference for carcasses. Accordingly, positive or negative values show preference or avoidance of carcass, respectively and thus negative values indicate preference for live prey.

The relationship between food preference, environmental factors, and explorative behavior was analyzed using generalized linear models (GLM). The initial full model contained Ivlev’s electivity index as dependent variable, and habitat complexity, turbidity, exploration rate, center use, and mobility rate as covariates. The interactions of environmental factors and exploration rate were also included. To find the most parsimonious model that includes all significant predictor variables, the backward elimination variable selection method was used (Grafen & Hails, 2002). To do this, the predictor variable with the highest p-value was removed from the full model, one after the other until the p-value of all remaining variables was below the threshold of *α*=0.05. Data were analyzed in the R 4.0.2 (R Core Team, 2020) statistical environment.

## Results

Black bullheads consumed food items in 22 out of 24 trials (92%). During the binary choice tests, 56% of carcass preys, and 36% of live preys was consumed. In five trials, black bullheads consumed all carcasses, but consumed live prey as well. We found that black bullheads consumed more carcass preys in complex habitat and clear water, whereas more live prey was consumed in simple habitat and turbid water (Fig. 2.). This relationship remained significant after model selection (Table 2., and Online Resource Table S1.). Exploration rate was generally high (mean ± SD: 61.88 ± 11.84 %), center use was low (2.10 ± 5.13 %) along with a high level of mobility rate (96.23 ± 4.12 %). We found no correlation of food choice and any measures of exploratory behavior (Online Resource Table S1.).

**Fig. 2.**
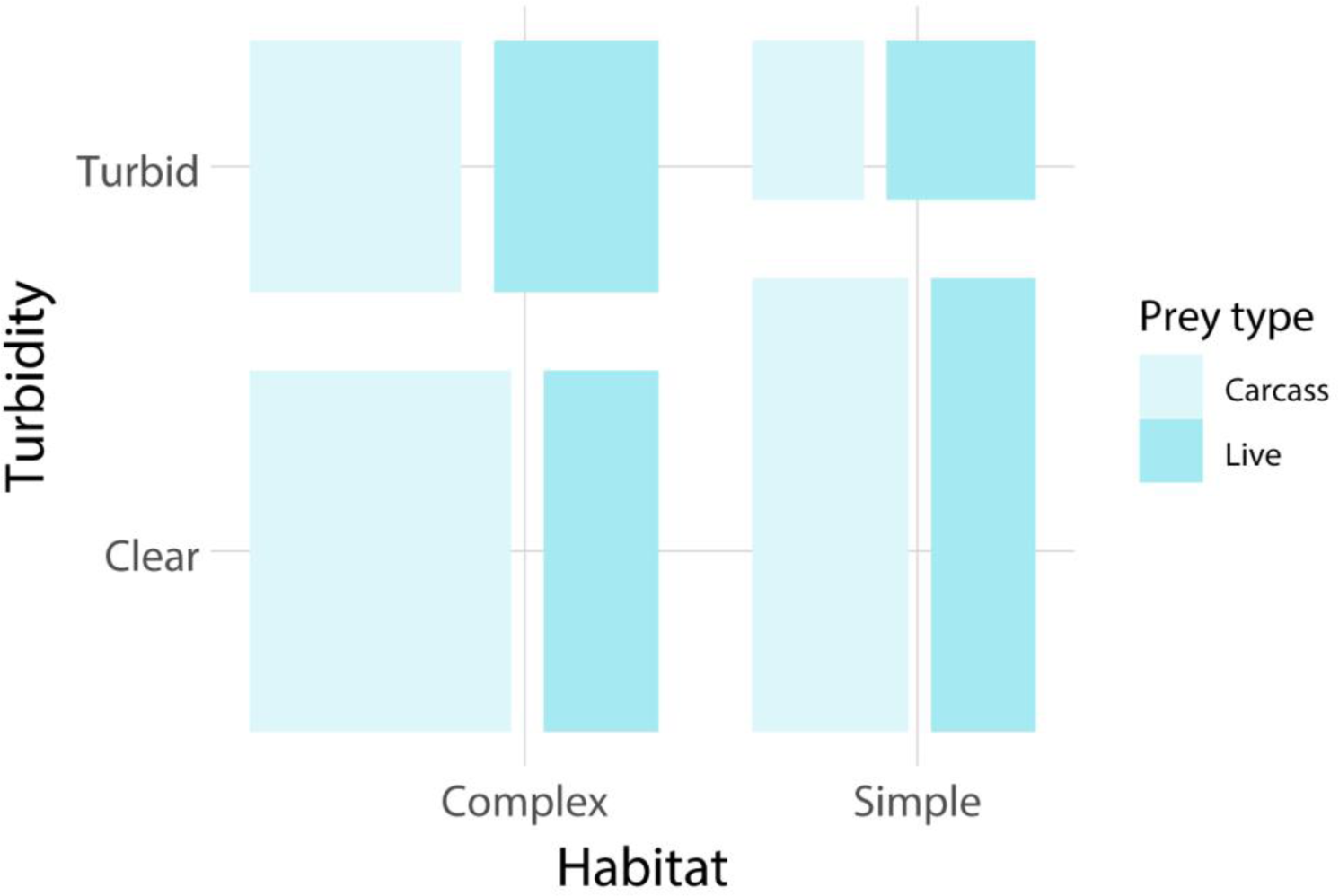
The consumption of carcass (N=40) and live prey (N=26) under different environmental setups. Size of the rectangles is proportional to the number of food items consumed (i.e. larger rectangle means more consumed food items)

**Table 2.**
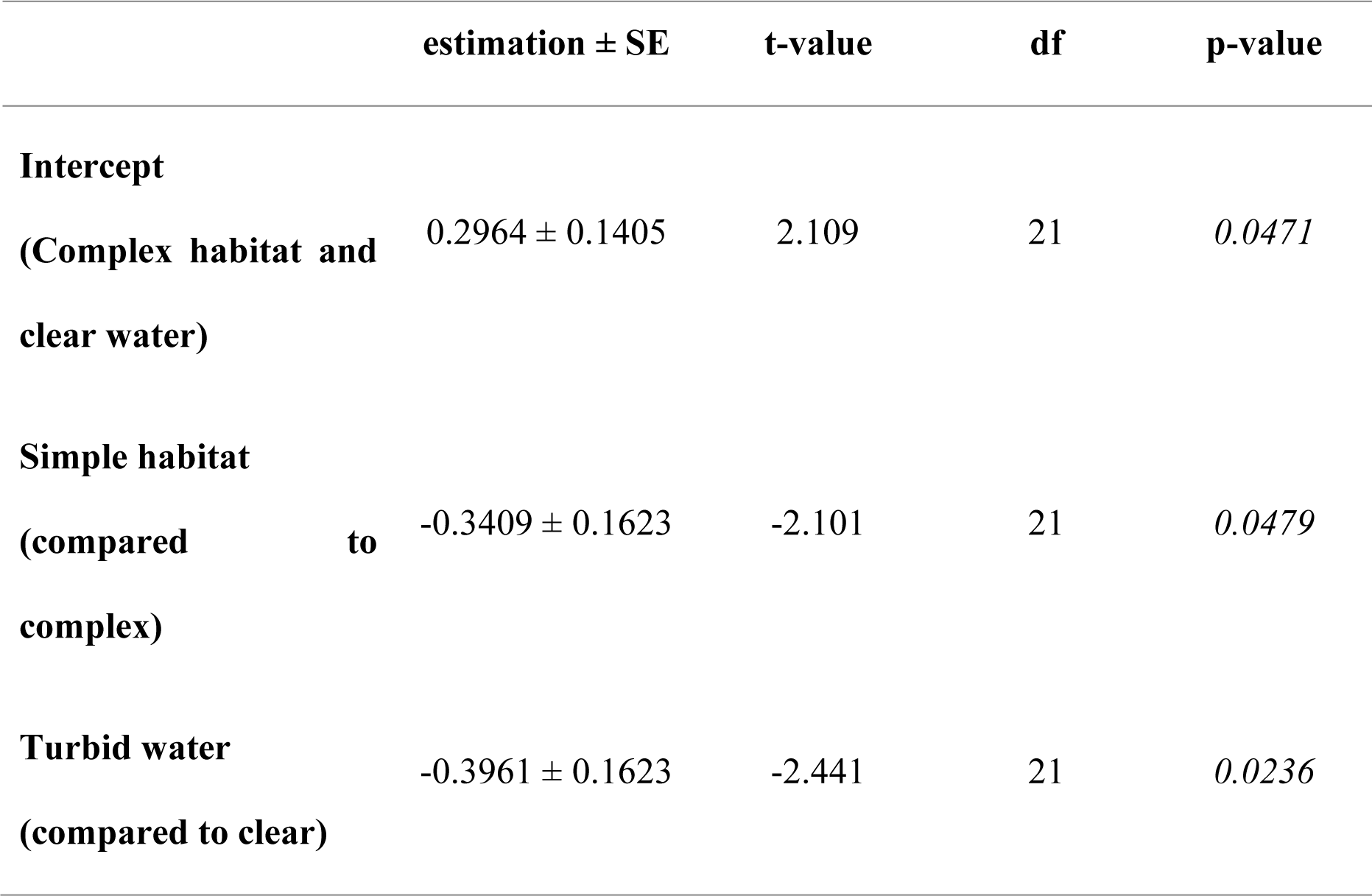
Results of the generalized linear model after model reduction. The relationship between environmental variables and Ivlev’s electivity index values of black bullheads (N=24). Positive values of the index indicate a preference for carcass prey, while negative values indicate a preference for live prey.

## Discussion

In this experiment we studied black bullheads’ foraging decisions under different environmental conditions. We offered them live and carcass preys of tubenose gobies in a binary food choice test simultaneously. We have also studied the relationship of these foraging decision and explorative behavior of the individuals. We demonstrated that black bullheads actively feed on live, fully mobile adult fish. Although we have found no correlation of explorative behavior and food choice, the latter is proved to be affected by turbidity and habitat complexity; black bullheads are more likely to choose carcass over live prey in clear water, or in more complex habitat.

Fish consumption of black bullheads has been confirmed in various habitats along the species’ geographical distribution, including its native and non-native range (see e.g. Leunda et al., 2008; and Snow et al., 2017). These studies although do suggest predation, they merely confirm the consumption of fish, but were not designed to distinguish between predation and scavenging, which is a documented way of fish consumption of black bullheads (Preiszner et al., 2020). In our study more than half of the black bullheads consumed live prey. Thus, to our knowledge, this experiment is the first to confirm predation on live adult fish by black bullheads. A species role in the food web of an ecosystem largely depends on its foraging tactics. The black bullhead is a successful invader in various freshwater ecosystems (Kottelat & Freyhof, 2007) and is capable of affecting native fauna (Jaćimović et al., 2023) not only via competition or interference (see e.g. Kreutzenberger, Leprieur, & Brosse, 2008) but if active predation is in its behavioral toolbox then by creating extra predation pressure on certain species’ populations (Leunda et al., 2008).

Among the various physiological effects of turbidity on fish, arguably a major consequence is the reduced transmission of light, and thus reduced visibility. Although prey may be more visible for predators in clear water, visual predator detection of prey is also more effective. Also, there is a contrasting effect in prey detection for predators; piscivores that detect their prey from a large distance may experience more adverse effects of turbidity than those detecting their prey from a short distance (reviewed by Palm, 2001). While visual perception does play a crucial role in schooling behavior for the members of the family Ictaluridae, where black bullheads belong, it is suspected to be inferior to other senses in foraging (Echelle, Kuhajda, & Ross, 2020). Ictalurids use chemo-orientation, likely to be regulated by gustation while searching for food (Daghfous, Green, Zielinski, & Dubuc, 2012; Kanwal & Finger, 1997; Parker, 1910). Gobies of the family Gobiidae on the other hand are suggested to use visual orientation extensively (Neiße, Santon, Bitton, & Michiels, 2020; Patzner, Van Tassell, Kovačić, & Kapoor, 2011), therefore in this case better visibility may have been more advantageous for the prey. Increased avoidance behavior of the gobies may have decreased the prey encounter rate of the predator. Hazelton & Grossman (2009) proposed that turbidity may facilitate invasive species’ effects on native populations through asymmetrically changing foraging success. As Palm (2001) pointed out, turbidity may change community structure in favor of species that are less reliant on visual orientation. A mechanism shaping this phenomenon may be that consumers which are better equipped with chemo- and mechanosensory apparatus could benefit more from flexible foraging tactics in turbid environments. Thus, such flexibility may play a role in black bullheads’ widespread invasion success, that is largely attributed to fishponds (Takács et al., 2017), which due to farming practices generally have elevated turbidity compared to surrounding natural habitats. This phenomenon may even have positive feedback for the species. For example Braig and Johnson (2003) demonstrated experimentally, that black bullheads may increase turbidity in some ecosystems.

Higher scavenging rate in complex habitat may be explained in various ways. Firstly, complexity of a habitat may alter behavior via modifying visibility; for example reducing visual encounter rate of prey by the predator. This may result decreased predation through a similar mechanism to turbidity, explained above. On the other hand, habitat complexity can benefit both prey and predator fish for example by active shelter use. Rocks, like submerged vegetation, and other structures, offer hiding places for various organisms. Black bullheads are not known as ambush predators however, thus a more complex habitat offering various options to utilize as shelters is probably more profitable for the gobies to incorporate in their predator evasion behavior. In Lake Balaton, where the individuals for this experiment were caught, tubenose gobies use rip-rap habitats extensively (Czeglédi et al., 2019) and the individuals caught for this experiment were exclusively from this habitat. Tubenose gobies’ use of rocks as shelters is a basic behavioral pattern. In case of better predator avoidance of tubenose gobies, consuming carcasses may have been more profitable for black bullheads than hunting for live prey. Thus, a feasible explanation of the experienced choice pattern is that it has emerged as a consequence of both the consumers’ and the preys’ behavior. Baber and Babbit (2004) found that same habitat complexity affects differently the predator’s foraging success on two prey species, and they argue that this is caused by prey behavior, namely the difference in their activity levels. Although in our experiment we used only one prey species, but in two distinct level of motility. Furthermore, piscivorous fish shift between prey species depending on turbidity levels, and also on cover availability for prey (Carter, 2010). Thus, a similar shift between carcass and live prey may be caused by turbidity and shelter availability.

Although we found no correlation between explorative behavior and food choice of black bullheads in our experiment, it may be important in other context, e.g. in presence of conspecifics or other competitors. For example, boldness did not covary with the speed of food choice of individually kept three-spined sticklebacks, *Gasterosteus aculeatus* (Linnaeus, 1758). In groups however, less bold individuals had increased exploration rate and decreased foraging speed, suggesting greater behavioral flexibility of these individuals (Ólafsdóttir & Magellan, 2016). Similar variation in flexibility may be experienced in black bullheads as they exhibit definitive shoaling behavior in early life stages. A further option is that our results are biased. Firstly, the length of the experiment combined with the size of the test aquaria could have masked the correlation between explorative behavior and food choice, potentially due to inherently high encounter rate. Secondly, although open field test is effective in estimating the personality trait of explorative behavior, it has it pitfalls and may be biased by other personality traits and current physiological state of the tested individuals (Gould et al., 2009; Perals, Griffin, Bartomeus, & Sol, 2017). The high rate of mobility despite the length of the open field test and the extensive wall-following behavior recorded in our experiment may suggest anxiety (Simon, Dupuis, & Costentin, 1994), and thus may have masked the actual individual propensity to explore.

In conclusion, we demonstrated experimentally that turbidity and habitat complexity may decrease the predation rate and increase scavenging behavior in black bullheads, which may be derived from both consumer and prey behavior. Understanding the relationship between environmental factors and foraging ecology is important for effective freshwater ecosystem management and conservation. The extent of the predatory behavior of black bullhead still has to be studied to fully understand the species’ ecological effect as an invader. Furthermore, to understand the relationship of personality and foraging decisions of scavenging fish, additional studies are encouraged. As elasmobranch fish are also known to be more susceptible for bait of anglers – which may be interpreted as scavenging – in more turbid circumstances (Ebert, 1991), it can be argued that a change in prevalence of scavenging behavior along environmental gradients is taxonomically pervasive, calling for further research in scavenger ecology of fish.

## Statements and Declarations

The authors have no competing interests to declare that are relevant to the content of this article.

## Ethical Statement

All procedures involving the handling and treatment of fish were in accordance with the permit for the delivery and use of aquatic animals for scientific purposes (permit reg. no.: VE-I-001/01890-3/2013, valid between 22 August 2013 and 21 August 2023, issued by the Food-Security and Animal Health Directorate, Governmental Office of Veszprém County, Hungary).

## Contributions

BP and TE conceived the study. All authors designed the methodology. BP, AS, and IC carried out the experiment. AS and BP analyzed the data, created the figures, and wrote the initial draft of the manuscript. All authors contributed critically to the drafts.

## Data Availability

The data that support the findings of this study are available from the corresponding author upon reasonable request.

## Online Resource

Table S1

Figure S1

Video S1

