## Supplemental Tables for "Effects of Turbidity and Habitat Complexity on the Foraging Behavior of the Black Bullhead (*Ameiurus melas*)"

### *on bioRxiv*

Bálint Preiszner[1,2*]; Anna Szolnoki[3], István Czeglédi[1,2], Tibor Erős[1,2]

[1] HUN-REN Balaton Limnological Research Institute, Fish and Conservation Ecology Research Group, Klebelsberg K. Street 3., 8237 Tihany, Hungary

[2] National Laboratory for Water Science and Water Security, HUN-REN Balaton Limnological Research Institute, Klebelsberg K. Street 3., 8237 Tihany, Hungary

[3] University of Veterinary Medicine, Institute of Biology, Department of Zoology, Rottenbiller street 50., 1077 Budapest, Hungary

**Table S1**

Results of the full generalized linear model. The relationship between variables and Ivlev's electivity index values of black bullheads (N=24). Positive values of the index indicate a preference for carcass prey, while negative values indicate a preference for live prey

| | estimation ± SE | t-value | df | p-value |
|---|---|---|---|---|
| **Constant (complex habitat, clear water)** | -0.003 ± 2.852 | -0.001 | 15 | 0.999 |
| **Normal habitat (compared to complex)** | 0.419 ± 1.204 | 0.348 | 15 | 0.733 |
| **Turbid water (compared to clear)** | 0.254 ± 1.040 | 0.244 | 15 | 0.811 |
| **Exploration rate** | 0.007 ± 0.012 | 0.549 | 15 | 0.591 |
| **Mobility rate** | -0.002 ± 0.027 | -0.085 | 15 | 0.934 |
| **Centre use** | -0.006 ± 0.020 | -0.317 | 15 | 0.755 |
| **Habitat × Turbidity** | -0.502 ± 0.379 | -1.326 | 15 | 0.205 |
| **Habitat × Exploration rate** | -0.009 ± 0.018 | -0.469 | 15 | 0.646 |
| **Turbidity × Exploration rate** | -0.007 ± 0.016 | -0.400 | 15 | 0.694 |

**Fig. S1**

Example of graphical outputs of the tracking software. Panel a) heat map (white represents unexplored unit, red represents the unit with the highest occurrence of the individual) of explored area of the arena, b) full track of the individual in the arena. The two examples are of different individuals

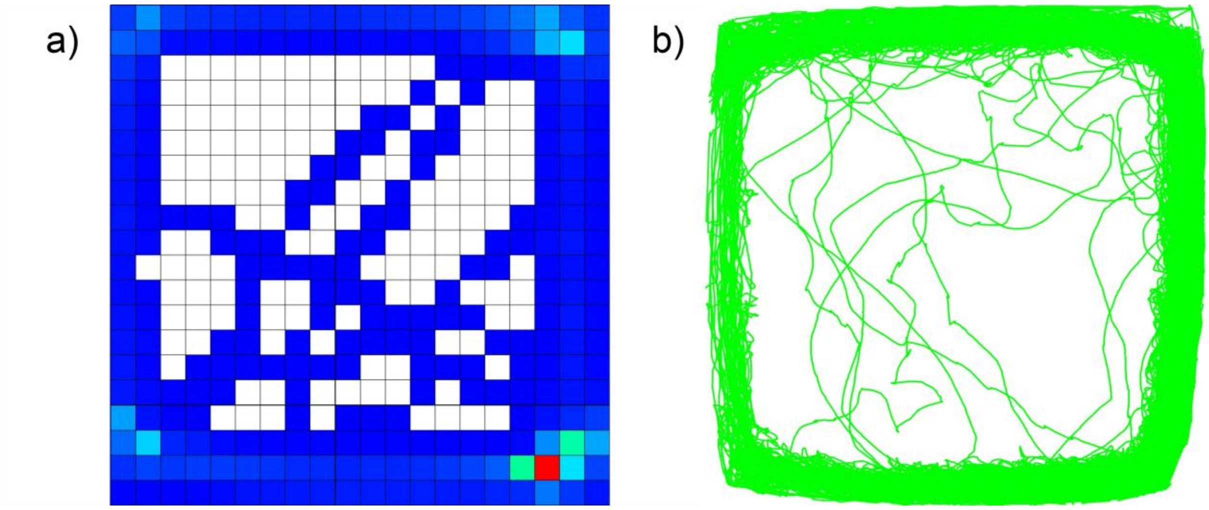


**Video S1**

Short example of open field test video. The footage shows an individual used during the pilot tests. The video is published along the online version of the article (Supplementary file 2)
